# The mechanosensitive ion channel Piezo1 contributes to ultrasound neuromodulation

**DOI:** 10.1101/2023.01.07.523089

**Authors:** Jiejun Zhu, Quanxiang Xian, Xuandi Hou, Kin Fung Wong, Tingting Zhu, Zihao Chen, Dongming He, Shashwati Kala, Jianing Jing, Yong Wu, Xinyi Zhao, Danni Li, Jinghui Guo, Zhihai Qiu, Lei Sun

**Affiliations:** Department of Biomedical Engineering, The Hong Kong Polytechnic University, Hong Kong SAR; Guangdong Institute of Intelligence Science and Technology, Hengqin, Zhuhai, Guangdong 519031, China; Guangdong Institute of Intelligence Science and Technology, Hengqin, Zhuhai, Guangdong 519031, China

**Keywords:** Transcranial ultrasound neuromodulation, Focused ultrasound, Mechanosensitive ion channels, Piezo1, Sonogenetics

## Abstract

Transcranial low-intensity ultrasound is a promising neuromodulation modality, with the advantages of non-invasiveness, deep penetration, and high spatiotemporal accuracy. However, the underlying biological mechanism of ultrasonic neuromodulation remains unclear, hindering the development of efficacious treatments. Here, the well-known Piezo1, was studied through a conditional knockout mouse model as a major molecule for ultrasound neuromodulation *ex vivo* and *in vivo*. We showed that Piezo1 knockout in the right motor cortex of mice significantly reduced ultrasound-induced neuronal calcium responses, limb movement and muscle EMG responses. We also detected higher Piezo1 in the central amygdala (CEA) which were found more sensitive to ultrasound stimulation than that of cortex. Knocking out the Piezo1 in CEA neurons showed a significant reduction of response under ultrasound stimulation while knocking out astrocytic Piezo1 showed no obvious changes in neuronal responses. Additionally, we excluded an auditory confound by monitoring auditory cortical activation and using smooth waveform ultrasound with randomized parameters to stimulate P1KO ipsilateral and contralateral regions of the same brain and recording evoked movement in the corresponding limb. Thus, we demonstrate that Piezo1 is functionally expressed in different brain regions, and that it is an important mediator of ultrasound neuromodulation in the brain, laying the ground for further mechanistic studies of ultrasound.

## Introduction

Ultrasound is an emerging technology capable of non-invasively modulating neurons in various brain regions, whether relatively deep or superficial, with minute spatial and temporal resolution. Multiple studies have demonstrated effective ultrasonic activation of the human cortical (Lee *et al*., 2015; Legon *et al*., 2014), sub-cortical (Legon, Ai, *et al*., 2018) and related networks (Lee *et al*., 2015; Lee, Kim, *et al*., 2016; Legon, Ai, *et al*., 2018; Legon, Bansal, *et al*., 2018; Legon *et al*., 2014). Basic clinical trials with ultrasound d have shown improvement in specific behavioral outcomes, such as improved mood (Reznik *et al*., 2020; Sanguinetti *et al*., 2020) and increased responsiveness in patients with chronic disorder of consciousness (Cain *et al*., 2021). Crucially, there were no observable side effects in these studies even with chronic stimulation. Multiple studies have also shown that ultrasound stimulation of specific neurons or brain regions can elicit distinct behaviors, such as motor or visuomotor response, in rodents (Kim *et al*., 2012; King *et al*., 2013; King *et al*., 2014; Tufail *et al*., 2010; Ye *et al*., 2016; Yoo *et al*., 2011; Yu *et al*., 2021), sheep (Lee, Lee, *et al*., 2016) and monkeys (Deffieux *et al*., 2013; Wattiez *et al*., 2017). Thus, a growing body of evidence shows ultrasound to be a promising neuromodulation modality – generally safe, deliverable non-invasively and apt for clinical translation. However, the biological mechanism underlying the neuromodulatory effects of ultrasound remains to be elucidated. This lack of clarity is a hurdle in the path of future ultrasound-based therapies if they are to be applied predictably and consistently, with maximal achievable efficacy and minimal side-effects.

Ultrasound comprises acoustic waves at frequencies above the upper limit of the audible range for humans (20 KHz). When exposed to ultrasound, nanometer-scale compression and particle displacement of the medium occurs, with heat, cavitation, or acoustic radiation force (ARF) generated when the particles return to their original configurations. Depending upon the energy level, ultrasound can be categorized as high- or low-intensity (Ter Haar, 2007). High-intensity ultrasound produces heat, and this has been used to develop high-intensity focused ultrasound (HIFU) (spatial-peak temporal average intensity, ISPTA > 1000 W/cm^2^). This has been applied as a treatment for several neurological diseases including essential tremor (Iorio-Morin *et al*., 2021), Parkinson’s disease (Sinai *et al*., 2022) and obsessive-compulsive disorder (Germann *et al*., 2021). However, ultrasound that can modulate neurons without cell damage would require low intensity (ISPTA < 500 mW/cm^2^), where the induced temperature change would be minor (<0.1°C) (Dalecki, 2004; Dinno *et al*., 1989; Kim *et al*., 2014; O’Brien Jr, 2007; Ter Haar, 2007; Tyler *et al*., 2018). Furthermore, mutant *C. elegans* worms with and without knockout of thermosensitive ion channels showed no difference in neuronal response to low-intensity ultrasound stimulation (Kubanek *et al*., 2018) implying a mechanical, rather than thermal, mechanism. While cavitation may be induced at higher intensities (Leighton, 2007), research using lower intensities showed that micron-scale tissue displacement-induced spiking activity remained unchanged within a broad range of acoustic frequency (0.5–43 MHz) (Menz *et al*., 2019) suggesting ARF as the dominant physical mechanism.

A pivotal component for cellular sensation of mechanical disturbances like ARF and enablers of fast neuromodulatory effects, mechanosensitive ion channels are plausible mediators of ultrasound (Fomenko *et al*., 2018; Tyler *et al*., 2018). Studies have shown ultrasound to be significantly more effective in cells overexpressing one of various such channels *in vitro* (Kubanek *et al*., 2016; Qiu *et al*., 2019; Sorum *et al*., 2021; Yoo *et al*., 2022), in a simple nervous system (Kubanek *et al*., 2018) and in animals (Qiu *et al*., 2020). Ultrasound was also found capable of indirect neuromodulation through activating astrocytic TRPA1 (Oh *et al*., 2019). These results suggest that the ultrasonic effect in the mammalian brain could be modulated by endogenous mechanosensitive ion channels that are present. Piezo1, the most sensitive mechanotransduction ion channel, is known to respond to force as low as 10 pN (Coste, 2010; Wu *et al*., 2016). Separately, hippocampal neurons have been shown capable of sensing localized mechanical force as low as 13-50 pN (Falleroni *et al*., 2022). Interestingly, online database (Lein *et al*., 2007; Sjöstedt *et al*., 2020) **show broad expression of Piezo1 RNA across both the human and mouse brain**. Taken together, these facts led us to hypothesize that Piezo1 could be mediating the neuromodulatory effects of ultrasound *in vivo*.

Our previous study showed the significant contribution of Piezo1 in the ultrasonic stimulation of neurons *in vitro* (Qiu *et al*., 2019). We, thus, wanted to investigate the role of Piezo1 in live animals in the present study. We first determined the functional expression of Piezo1 in the mouse brain with the generation of Piezo1 knockout (P1KO) neurons. The functional expression of Piezo1 and the success of conditional knockout was confirmed by genotyping, immunostaining and calcium imaging with the Piezo1 agonist Yoda1. We then tested the neuronal activity induced by ultrasound both *ex vivo* and *in vivo*. In acute brain slices, ultrasound-induced calcium influx was significantly reduced in P1KO neurons and in neurons pre-treated with the broad mechanosensitive ion channel blocker ruthenium red (RR) (Drew, 2002) compared to those with functional Piezo1. We also used *in vivo* motor responses as a gauge of ultrasound stimulation as in previous studies (Aurup *et al*., 2021; Ye *et al*., 2016). P1KO mice displayed lowered responses to ultrasound stimuli in the motor cortex, measured by reduced contralateral limb movement, reduced muscle EMG responses, and reduced calcium signaling and c-Fos expression. Further, in a brain region showing higher Piezo1 expression than the cortex, neurons showed greater sensitivity to ultrasound stimulation as well. Taken together, our loss-of-function and agonist-stimulated experiments demonstrate Piezo1 as a major mediator of ultrasonic neuromodulation *in vivo*. We also excluded an auditory confound by monitoring c-Fos expression in the auditory cortices of control, P1KO and deafened mice. In short, the present study demonstrates that Piezo1 is a key mediator for ultrasonic neuromodulation in the mouse brain and provides a basic study of its spatial distribution in different brain regions.

## Results

### Piezo1 mediates neuronal responses to ultrasound stimuli in *ex vivo* brain slices

We first examined whether Piezo1 was expressed endogenously at a sufficiently-high level to be functionally detected. Previous studies have used neurons of the mouse cortex for *in vitro* studies (Qiu *et al*., 2019; Yoo *et al*., 2022) as well as for *in vivo* motor cortex stimulation (King *et al*., 2014; Tufail *et al*., 2010; Ye *et al*., 2016). Hence, we opted to build upon this literature by testing Piezo1’s possible contribution to ultrasound neuromodulation in the mouse cortex. We found cortical cells co-expressing Piezo1 and MAP2 (Fig 1A) confirming the expression of Piezo1 in cortical neurons. The Piezo1 antibody’s specificity was verified by confirming a loss of observed staining when brain tissue was pre-incubated with the antibody’s specific blocking peptide (Fig S1). We also found neurons co-stained for GFAP (an astrocytic marker) and PIEZO1. indicating the expression of Piezo1 in astrocyte as well (Fig S2).

**Fig 1.**
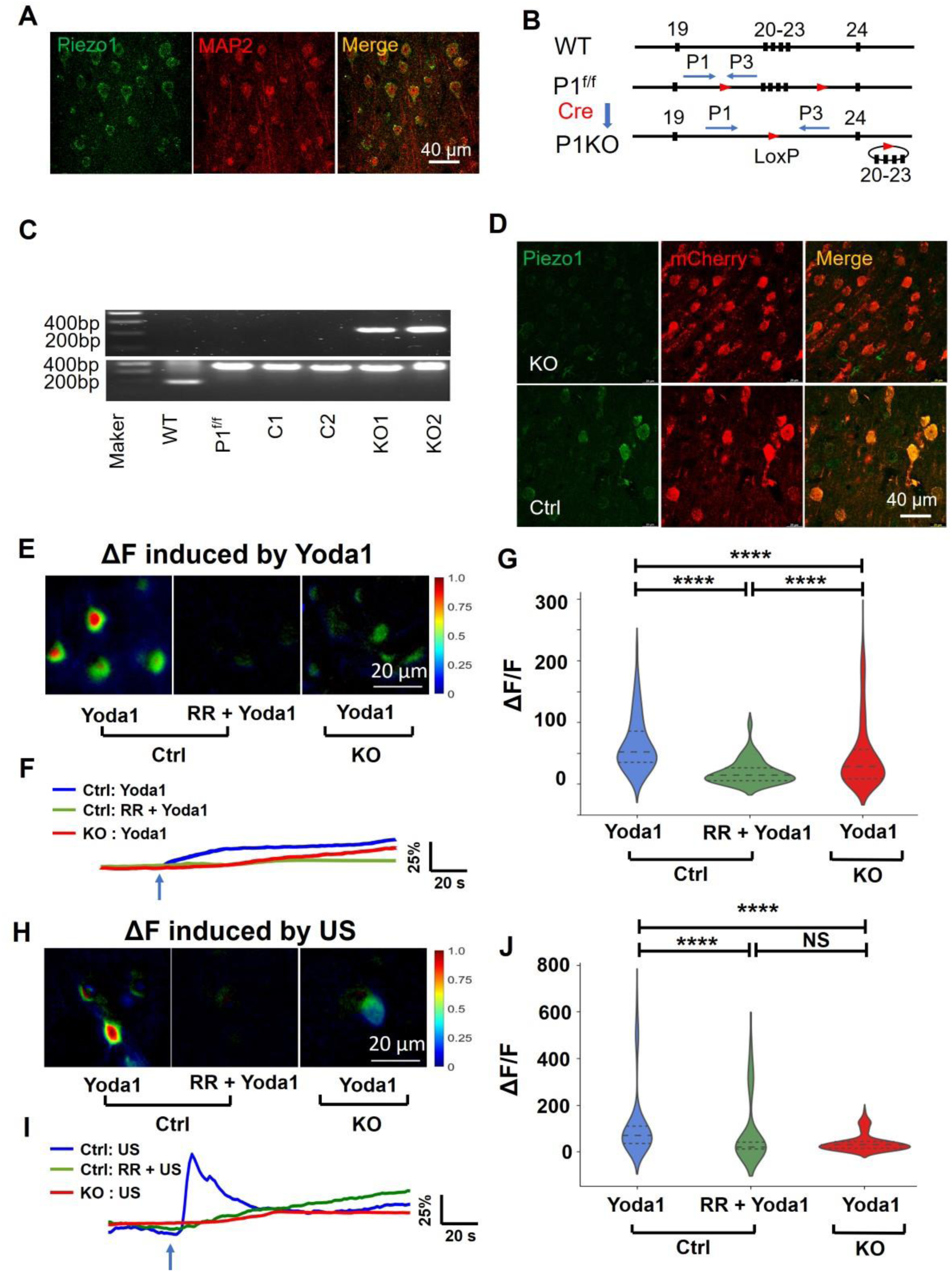
Piezo1 contributes to neuronal responses to ultrasound stimuli in *ex vivo* brain slices. **(A)** Immunostaining of Piezo1 and MAP2 of the cortex of WT mice. **(B)** Schematic illustration of Piezo1 conditional knockout by Cre-LoxP system. **(C)** Top: genotyping results with deletion-spanning PCR primers. Expected PCR product size is 230 bp after deletion and 3080 bp without deletion which cannot be amplified; Bottom: genotyping results with LoxP-spanning PCR primers. Expected size in Piezo1^f/f^ mice = 380bp, WT = 188bp. **(D)** Immunostaining of Piezo1 in Ctrl and P1KO mice brains. **(E)** Representative images of *ex vivo* neuronal calcium signaling from brain slices of Ctrl or P1KO mice after adding 20 μM Yoda1. **(F)** Representative calcium traces of one neuron from brain slices of Ctrl, Ctrl+RR or P1KO mice after adding 20 μM Yoda1. **(G)** Summarized calcium changes of neurons from *ex vivo* experiments. n is as follows: 208 neurons from 4 Ctrl mice; 66 neurons from 4 Ctrl mice + RR; 316 neurons from 5 P1KO mice. Bars represent the medium and interquartile range ****p < 0.0001, by one-way ANOVA with post-hoc Dunn’s test. **(H)** Representative images of *ex vivo* neuronal calcium signaling from brain slices of Ctrl or P1KO mice treated with 0.35Mpa US. **(I)** Representative calcium traces of one neuron from brain slices of Ctrl, Ctrl+RR or P1KO mice treated with 0.35Mpa US stimulation. **(J)** Summarized calcium changes from *ex vivo* experiments. n is as follows: 277 neurons from 4 Ctrl mice; 98 neurons from 4 Ctrl mouse + RR; 283 neurons from 5 KO mice. Bars represent the medium and interquartile range, ****p < 0.0001, ‘N.S.’ not significant, by one-way ANOVA with post-hoc Dunn’s test.

To evaluate the role of Piezo1 in ultrasound neuromodulation, mice with conditional knockout of Piezo1 in neurons of the cortex were generated using the Cre-LoxP system (Fig 1B). Mice with exons 20–23 of Piezo1 flanked by loxP sites (Cahalan *et al*., 2015) were bred, and Cre recombinase-mCherry virus was delivered to the cortex by local injection, thus generating mice with Piezo1 knockout (P1KO) in cortical neurons (Fig 1B). A virus coding for mCherry alone was used as a control (Ctrl). PCR genotyping with primers spanning the deletion showed the expected pattern (Cahalan *et al*., 2015), with a 230 bp band seen exclusively in P1KO mice (Fig 1C), indicating successful a Piezo1 knockout model. Immunostaining for Piezo1 showed much reduced fluorescent signals in P1KO neurons compared to Ctrl (Fig 1D, Vid S1), confirming the knockout of Piezo1 in neurons of the targeted region.

The functioning of endogenous Piezo1 was then tested by calcium imaging, using a pharmacological agonist or antagonist, and ultrasound stimulation. An additional virus encoding the genetically-encoded calcium sensor GCaMP6s was delivered along with Cre-mCherry or mCherry viruses (Fig S3A). This generated P1KO and Ctrl mice with GCaMP6s neuronal expression in their motor cortices. Acute brain slices were collected four weeks post-injection and treated with Yoda1 or ultrasound, in a setup building by a 0.5 MHz plane transducer, a gassed perfusion system and inverted fluorescent microscope (Fig S3B-C). Calcium influxes were observed when Ctrl brain slices were treated with 20 μM of the Piezo1 agonist Yoda1 (ΔF/F= 60.36% ± 43.92%), but this was significantly reduced in the RR+Yoda1 condition (14.27% ± 17.26%), and in P1KO neurons treated with Yoda1 (46.88% ± 54.95%) (Fig 1E-G). Neurons in the acute brain slices were then treated with the pulses of ultrasound (US) at 0.35 MPa with 1kHz PRF, 50% duty cycle, for 300 ms, to test their ability to respond to ultrasound stimuli. Consistent with the Yoda1 pattern, Ctrl neurons showed significantly stronger calcium response to ultrasound sonication (115.3% ± 147.8%) than Ctrl + RR (83.83% ± 146.2%) or P1KO neurons (38.65% ± 38.27%) ((Fig 1H-J), and there is no significant difference between the latter two groups’ responses. These imaging data confirm the functional expression of Piezo1 channels in the given regions, as well as the neurons’ ability to respond to the mechanical stimulus of ultrasound and the reduced Piezo1 expression in P1KO mice. Taken together, we found that Piezo1 was functionally expressed in cortical neurons of mouse brains, and it was an important mediator for ultrasound neuromodulation in *ex vivo* brain slices.

### Cortical Piezo1 is an important mediator for ultrasound-induced motor behavior *in vivo*

We further studied evoked motor responses to evaluate the role of Piezo1 in ultrasound neuromodulation at the behavioral level. In P1KO or Ctrl mice, ultrasound stimuli of varying intensities were delivered to the right motor cortex, and responses in the left fore- and hind-limb were monitored using a high-speed camera and EMG measurement (Fig 2A, B). Ultrasound applied was of 0.5 MHz frequency, 1kHz PRF, 50% duty cycle. Consistent with previous studies, contralateral limb movement and EMG signal could be induced by ultrasound stimulation (Ye, *et al*., 2016; Aurup *et al*., 2021) (Vid S2). In both Ctrl and P1KO mice, forelimb movement was found to increase with the pulse duration under 0.35 MPa ultrasound (Fig 2C). However, the movement distance in P1KO was significantly lower than Ctrl mice at 250 ms (0.83 ± 0.74 mm vs. 3.78 ± 1.40 mm) and 500 ms (1.25 ± 1.12 mm vs. 4.84 ± 0.44 mm) pulse duration (Fig 2C). The hindlimb showed a similar pattern but showed a better response to a higher ultrasound intensity, 0.45MPa (Fig 2C). Nevertheless, the movement induced under these parameters was significantly lower in P1KO mice than in Ctrl mice (1.47 ± 0.90 mm vs 3.59 ± 1.72 mm.; Fig 2C). EMG responses in P1KO mice were obviously diminished, although not abrogated, compared to the Ctrl (Fig 2E). Upon quantifying EMG amplitude, we found that the response in both Ctrl and P1KO mice increased with growing ultrasound intensities, and the responses in P1KO mice were lower than the Ctrl at every intensity tested (Fig 2F).

**Fig 2.**
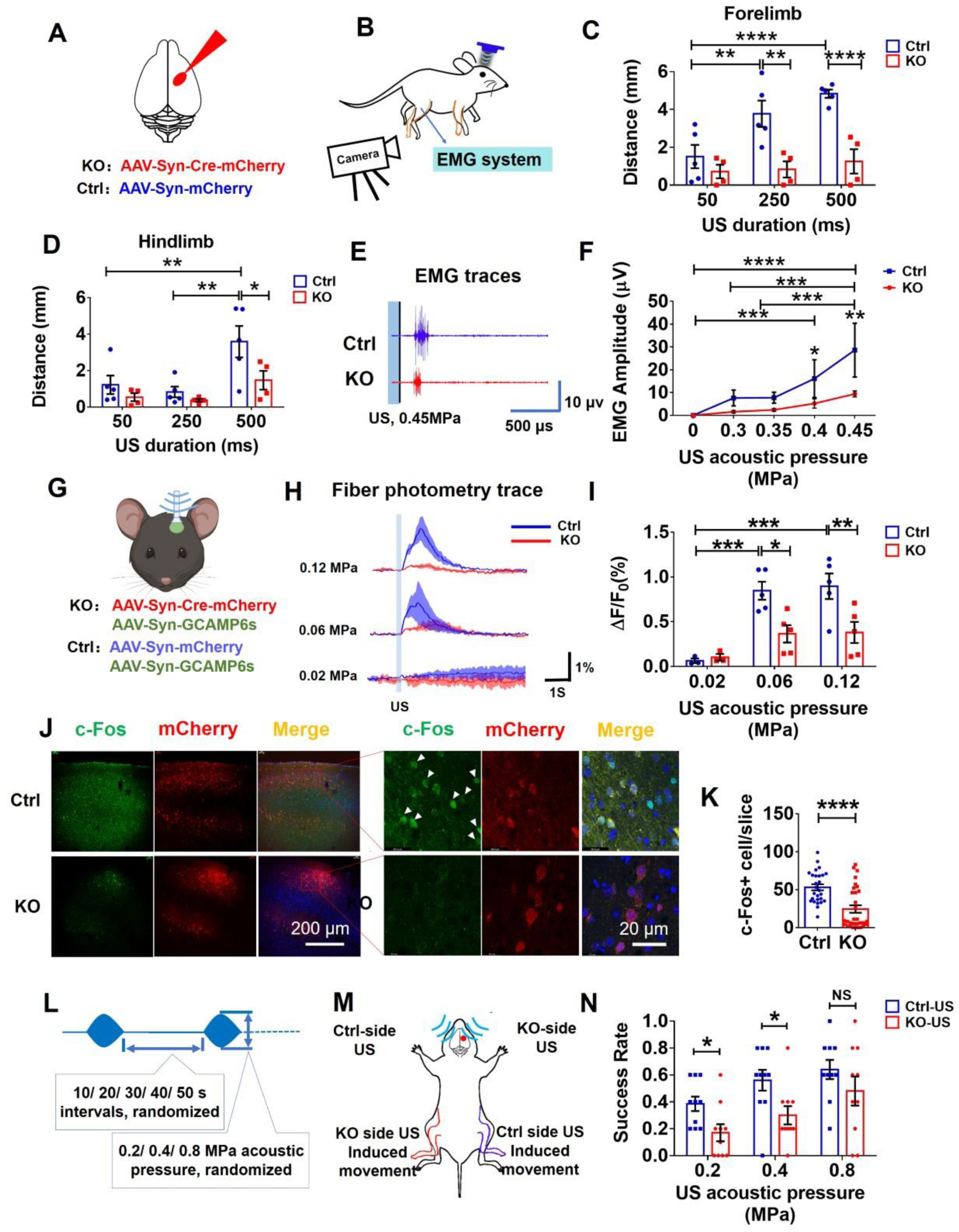
Cortical Piezo1 is an important mediator for ultrasound-induced motor behavior *in vivo*. **(A)** Schematic illustration of virus injections into mouse cortex for obtaining Ctrl and P1KO mice. **(B)** The right motor cortex of Ctrl or KO mice was treated with ultrasound stimuli at different parameters, and responses in the left forelimb and hind limb were monitored with a high-speed camera and EMG measurement. **(C)** Summarized forelimb movement upon ultrasound stimulation. N = 5 for Ctrl and 4 for P1KO mice, Bars represent mean ± SEM, *p < 0.05 and **p < 0.01 by two-way ANOVA followed by Sidak’s test. Ultrasound parameters were: Frequency: 0.5MHz, PRF: 1kHz; Amplitude: 0.35 MPa, Duty cycle: 50%; 5 s intervals, 6-10 times repetition for each US duration. **(D)** Summarized hindlimb movement at varying ultrasound stimulation durations. N = 5 for Ctrl and 4 for P1KO mice. Bars represent mean ± SEM, *p < 0.05 by two-way ANOVA followed with post-hoc Sidak’ test for group factor and Tukey test for duration factor. Ultrasound parameters were Frequency: 0.5MHz, PRF: 1kHz, Amplitude: 0.45 MPa; Duty cycle: 50%; 5 s intervals, 6-10 times repetition in each US duration. **(E)** Representative EMG traces induced by ultrasound stimulation in Ctrl (top) and P1KO (bottom) mice. **(F)** Summarized EMG result at varying acoustic pressures. The ultrasound parameters were, Frequency: 0.5MHz, Duty cycle: 50%, Duration: 500ms, 15 s intervals, 4-6 times repetition for each group. n = 5 for Ctrl and 4 for P1KO mice, Bars represent the mean ± SEM, *p < 0.05, **p < 0.01, ***p < 0.001 and ****p < 0.0001 by two-way ANOVA followed by Tukey’s test. **(G)** Schematic illustration of virus injections for fiber photometry in Ctrl and KO mice. **(H)** Representative calcium signaling traces induced by ultrasound stimulation in Ctrl (top) and P1KO (bottom) mice at different acoustic intensities. **(I)** Summarized calcium signals induced by ultrasound stimulation in 5 Ctrl mice and 5 P1KO mice. Bars represent mean ± SEM; *p < 0.05, **p < 0.01 and ***p < 0.001 by two-way ANOVA with post-hoc Sidak test. **(J)** Immunostaining of c-Fos in motor cortex of Ctrl and P1KO mice after 0.45 MPa US stimulation for 50 mins. Left: low magnification images; Right: zoomed-in images at high magnification. Blue: DAPI, Green: c-Fos, Red: m-Cherry. **(K)** c-Fos^+^ cells in 27 slices from 3 Ctrl mice, and 31 slices from 3 P1KO mice. Bars represent mean ± SEM, ****p < 0.0001 by unpaired t test. **(L)** Schematic illustration of ultrasound stimulation parameter for motor behavioral test. Smoothed-waveform ultrasound was used, amplitude: 0.2/0.4/0.8 MPa; duty cycle: 50%; PRF: 1 kHz; duration: 500 ms; pulse interval: 10/20/30/40/50s. Every pulse as triggered randomly. **(M)** Schematic illustration of motor behavioral test with ultrasound stimulation either on the virus injection side or the contralateral side. Limb movement in response to each US pulse delivered was recorded using a camera **(N)** Summarized mouse hindlimb movement at varying acoustic intensities. N = 5, bars represent the mean ± SEM, *p < 0.05 by two-way ANOVA followed by Sidak’s test. Ultrasound parameters were: frequency: 0.5MHz; PRF:1 kHz; amplitude: 0.2/0.4/0.8 MPa; duty cycle: 50%; duration: 500ms; each pulse was triggered randomly at 10/20/30/40/50s intervals.

To directly measure neuronal activity, we used fiber photometry to record calcium signal changes in P1KO and Ctrl mice expressing GCaMP6s (Fig 2G, S4). Ultrasound at 0.5 MHz frequency, 1kHz PRF, 50% duty cycle, 300ms duration and acoustic pressures between 0.02 – 0.12 MPa were used to stimulate the brains. We found that ultrasound rapidly and transiently activated neurons, as measured by GCaMP6s fluorescence change, at all tested acoustic pressures, and this response was also found to increase with acoustic pressure (Fig 2H). However, responses from P1KO cells were significantly lower than the Ctrl (Fig 2H-I).

Aside from real-time responses, the expression of the activation marker c-Fos was also evaluated as a post-stimulation measure of neuronal activity (Fig 2J). Compared to Ctrl mice, P1KO mice showed significantly reduced c-Fos levels (24.32 ± 27.00 vs. 53.04 ± 20.40; Fig 2J-K). We also monitored c-Fos expression in auditory cortices of these mice to evaluate the known auditory confound effect, known to be a possibility with ultrasonic neuromodulation (Guo *et al*., 2018) (Sato *et al*., 2018)). No obvious differences was found between Ctrl and P1KO mice (90.36 ± 17.41 vs. 93.36 ± 34.21, Fig S5), indicating a low chance of auditory effects being the cause of the observed behavioral differences.

To further exclude the possible auditory confound, we perform ultrasound stimulation on P1KO site and their contralateral site by using smoothed waveforms while observing the contralateral responses (Mohammadjavadi *et al*., 2019). The ultrasound stimulation is trigged randomly to diminish the accumulated effect by the previous stimulations (Fig 2L). 5 P1KO mice is then prepared as before and treated with the ultrasound stimulation either the ipsilateral hemisphere (where virus is injected, KO side) or the contralateral (contralateral side of virus injection, Ctrl side) (Fig 2M). The success rate of the contralateral limb movement under each ultrasound pulses were measured. We found that the KO side ultrasound stimulation showed a lower successes rate inducing the limb movement in mice especially under the 0.2 MPa (0.37 ± 0.16 vs. 0.18 ± 0.18) and 0.4MPa (0.54 ± 0.23 vs. 0.27 ± 0.21) ultrasound stimulation (Fig 3N). And this phenomenon is dismissed when ultrasound intensity reached 0.8MPa which may indicate an alternative mechanism under higher intensity ultrasound neuromodulation. These results confirms that piezo1 contributes to ultrasound neuromodulation specifically rather than auditory confound.

### Neuronal Sensitivity to ultrasound stimuli is dependent upon the presence of neuronal Piezo1

Having found endogenous Piezo1 to be an important contributor to ultrasound neuromodulation in the motor cortex, we were led to wonder whether other brain regions, possibly expressing greater levels of Piezo1, may also serve as effective targets of ultrasound. We performed immunofluorescent staining of WT mouse brains and found several brain areas with strong Piezo1 expression, including the central amygdala (CEA), bed nucleus of the stria terminalis (BNST), paraventricular nucleus of hypothalamus (PVN), Edinger-Westphal nucleus (EW) and red nucleus (RN) (Fig 3A, Fig S6). The levels of Piezo1 expression in these brain regions were quantified by their fluorescent intensities and were all found to be greater than in the cortex (Fig 3B). Of these, we were interested in the CEA as a classic and well-established target of neuroscientific experiments controlling defensive and appetitive behaviors (Fadok *et al*., 2018). To see if Piezo1 is an important mediator, either in a direct or indirect way, mediating the ultrasound neuromodulation in this brain area, fiber photometry experiment is conducted. Piezo1 expression was knocked out in either neurons or astrocytes of the CEA by viral delivery, and all groups were made to express GcAMP6s (Fig 3C). This enabled the creation of 4 groups, named as follows: ‘hSyn-Cre’ (neuronal P1KO); ‘hSyn’ (neuronal viral Ctrl); ‘GFAP-Cre’ (astrocytic P1KO); ‘GFAP’ (astrocytic viral Ctrl). Neuronal calcium dynamics in response to ultrasound stimuli were then monitored through fiber photometry in all four groups (Fig S7). We found a clear, rapid, and consistent increase in GCaMP6s fluorescence to ultrasound pulses of very low intensity (0.12 MPa, 300 ms pulse duration, 5 s interval) in the hSyn, GFAP and GFAP-Cre groups but little-to-no response in the hSyn-Cre group (Fig 3D). We did not observe any repetitive patterns of fluorescence changes prior to the US stimulation. The ultrasound parameters used did induce a small Ca^2+^ response in the hSyn-Cre group (0.12 MPa, peak ΔF/F_0_ = 0.41% ± 0.077%) but the peak amplitude in other three groups was significantly higher at same ultrasound intensity (hSyn, ΔF/F_0_ = 1.13% ± 0.17%; GFAP-Cre, ΔF/F_0_ = 1.05% ± 0.12%; GFAP, ΔF/F_0_ = 1.12% ± 0.10%) (Fig 3D, 3E). Notably, the peak amplitude was found to increase as the applied ultrasound acoustic pressure increased from 0.03 MPa to 0.25 MPa, indicating that the neuronal Ca^2+^ responses were dependent on the amount of ultrasound energy applied (Fig 3F). In all tested cases, the hSyn-Cre group showed significantly lower calcium responses, indicating the importance of neuronal Piezo1 for ultrasound neuromodulation. Since Piezo1 levels were found to higher in the CEA than the cortex, we further compared the neuronal responses to ultrasound stimulation between these regions. Quantification of the ∆F/F peaks revealed that the slope of the ∆F/F peak-pressure relation in CEA neurons (k = 4.885) was greater than that of the cortex (k = 1.132) (Fig 3G). Thus, we found that CEA neurons displayed higher Piezo1 expression as well as higher sensitivity to ultrasound compared to the cortex, with acoustic pressures as low as 0.06 – 0.25 MPa. The tunability of the responses to ultrasound parameters also helped to confirm that the observed responses were triggered by ultrasound, and not background signals. These results further confirm that Piezo1 is an important factor mediating ultrasonic neuron modulation *in vivo*.

**Fig 3.**
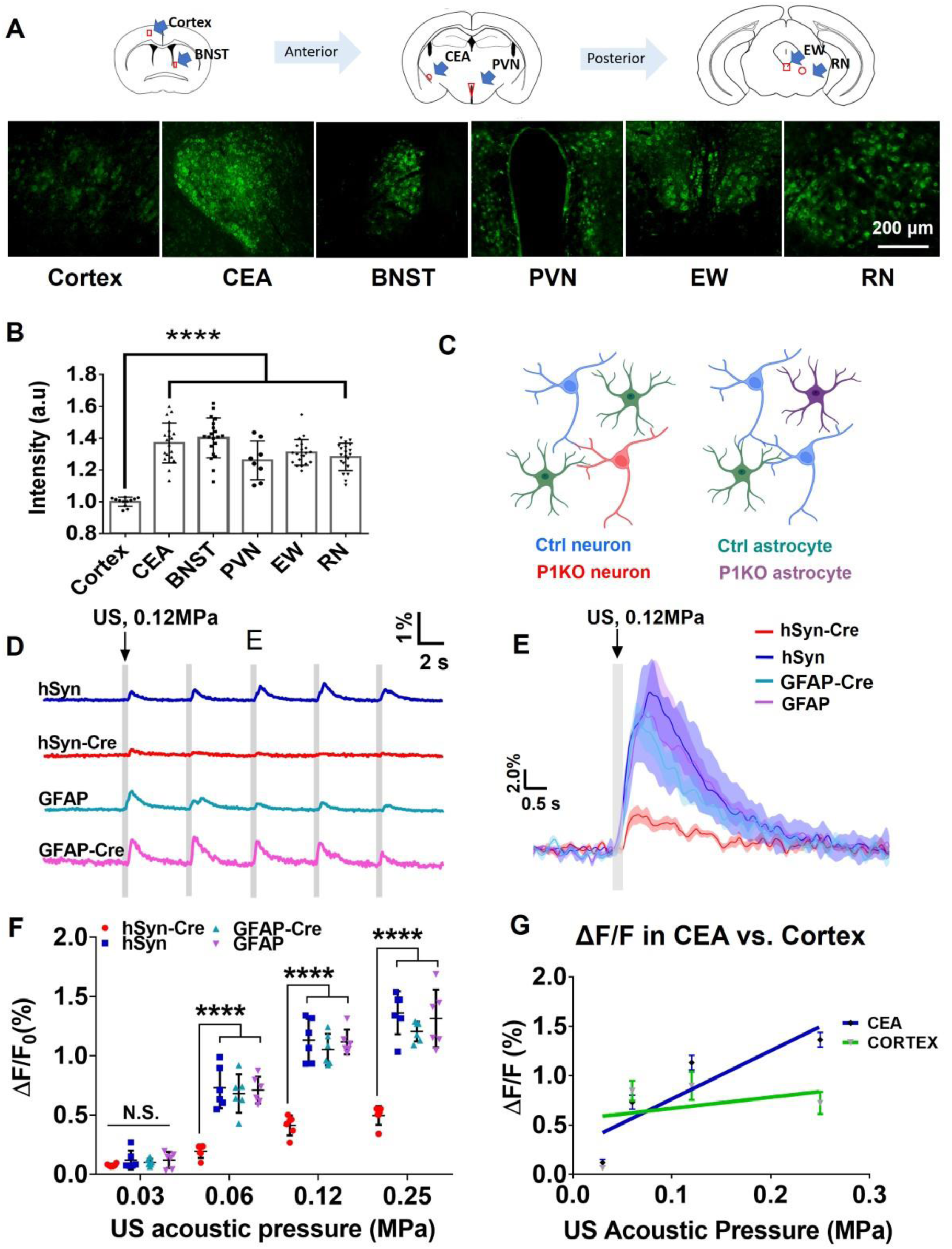
Neuronal sensitivity to ultrasound stimulation depends on the expression of neuronal Piezo1. **(A)** Immunostaining of Piezo1 in different brain regions, including the cortex, CEA, BNST, PVN, EW and RN. **(B)** Summarized relative fluorescence intensity of Piezo1 positive cells in different brain areas compared to that in cortex. Bars represent mean ± SD, ****p < 0.0001 by one-way ANOVA followed by Dunnett’s multiple comparisons test. **(C)** Schematic illustration of viral treatment in mice to create 4 groups for fiber photometry, with neuronal or astrocytic Piezo1 knockout. All groups were injected with hSyn-GcaMP6s **(D)** Representative calcium response traces under 0.12MPa ultrasound stimulation in the CEA. **(E)** Representative averaged calcium response traces following one ultrasound stimulus of 0.12MPa. **(F)** Summarized calcium response to varying ultrasound stimulations. N= 6 mice per viral group. Bars represent mean ± SEM, *p < 0.05, ***p < 0.001, ****p < 0.0001 by one-way ANOVA with post-hoc Tukey test. Ultrasound parameters were: Frequency: 0.5MHz; Duty cycle: 50%, PRF: 1 kHz, Duration: 300 ms; 5 s intervals; 5 times repetition in each intensity groups. **(G)** Regression of calcium signal induced by ultrasound in the cortex and CEA from 6 mice and 5 mice respectively. CEA:y=4.885x + 0.2743; Cortex: y=1.132x + 0.5542; Slope difference: p=0.003388.

## Discussion

In this study, we demonstrate that Piezo1 is an important mediator of the ultrasonic neuromodulatory effect to regulate neuronal signaling and animal motor behavior *in vivo*. We first confirmed the functional expression of Piezo1 in the mouse brain through the generated P1KO mouse model. We found that P1KO neurons displayed lower neuronal activity responding to ultrasound through the reduced ultrasonic motor response, EMG signaling, calcium changes and c-Fos expression compared to the control. A similar phenomenon was also found in CEA region; with the knockout of neuronal Piezo1 in the CEA, neurons showed significantly reduced ability to respond to ultrasound. In contrast, CEA neurons, having higher Piezo1 expression levels, displayed a greater sensitivity to ultrasound than cortical neurons, showing stronger responses at lowered acoustic pressures. We also addressed the auditory confound question by monitoring auditory cortex activation in P1KO and Ctrl mice, as well as applying smoothed waveform ultrasound. Thus, we used a dual approach of comparing neuronal responses to US in the same brain region with Piezo1 knocked out (P1KO vs Ctrl), and different brain regions with varying Piezo1 levels (CEA vs Cortex), to demonstrate the contribution of Piezo1 as mediator of ultrasonic neuromodulation *in vivo*. Additionally, although we found Piezo1 expressed in both neurons and astrocytes, we found that neuronal Piezo1 played a major role in mediating ultrasound’s effects directly.

While previous studies have identified various mechanosensitive ion channels which respond to ultrasound stimuli *in vitro*, studies with direct observation *in vivo* remain limited. It bears repeating that a transgenic animal model with conditional knockout of the ion channel Piezo1 was utilized in this study, allowing precise manipulation of its presence or absence to examine its specific role. We also demonstrated increased response to ultrasound in a region with greater Piezo1 expression. *In vivo* studies, naturally, provide more realistic scenarios in terms of the cellular/subcellular expression profile of different mechanosensitive ion channels which likely affect their distinct role within a brain network under ultrasound stimulus. We believe that our investigation with a verified KO system, and comparing neurons with differing endogenous levels of Piezo1, *in vivo* is useful for understanding the mediator roles relevant to ultrasound neuromodulation. These findings add to the literature by demonstrating the potency of ultrasound neuromodulation in the brain even when only endogenous levels of mechanosensitive proteins are involved.

Although the high sensitivity of Piezo1 to sense the mechanical cues (Coste, 2010) may explain its ability to modulate neurons, we also expect that there may be an amplification effect through cooperation with other components. For instance, TRPV4 or SMPD3 may contribute to this Piezo1 mediated ultrasonic neuromodulatory effects through their enhancement in Piezo1 function (Shi *et al*., 2020; Swain & Liddle, 2021). However, there is no direct evidence demonstrating this hypothesis. We believe further studies that focus on Piezo1’s cooperation with other participants in mechanotransduction under ultrasound stimulation may provide more information about the mechanism.

For the moment, we cannot exclude the role of other cellular machinery since ultrasound neuromodulatory effects were not fully eliminated in the P1KO neurons. Previous studies have shown that some TRP channels such as TRPA1, TRPP1/2, TRPC1, TRPV1 (Burks *et al*., 2019; Oh *et al*., 2019; Yang *et al*., 2021; Yoo *et al*., 2022), potassium channel such as TREK1 and TRAAK (Lengyel *et al*., 2021; Sorum *et al*., 2021), and calcium voltage-gated channels (VGCs) (Arnadottir & Chalfie, 2010; Tyler, 2012), may also contribute to the ultrasound induced neural activity. We agree with these findings that other ion channels in difference cell type, astrocyte(Oh *et al*., 2019) for example, may contribute to the ultrasound neuromodulation, especially under high ultrasound acoustic pressure. In the present study, we were able to use low acoustic intensities due to Piezo1’s highly mechanosensitive nature. However, at higher intensities it is understood that other ion channels or mechanotransduction elements, less mechanosensitive than Piezo1, may take over as a co-player mediating neuromodulation, thus diluting the role of Piezo1. Hence, we believe that under low ultrasound intensity, Piezo1 in neurons are the major mediator mediating the ultrasound neuromodulation *in vivo*.

An interesting finding slightly orthogonal to the purpose of this study was that we found Piezo1 highly expressed in the BNST, CEA, PVN, EW and RN. These are all corticotropin releasing factor (CRF)-expressing areas that regulate stress processing in animals (Henckens *et al*., 2016; Smith & Vale, 2022). CRF is a 41 amino acid peptide which is relatively large compared to other neural transmitters. Neurons might experience more mechanical effects while transporting and secreting this neuron transmitter (Tyler, 2012) when processing stress in animals thus requires more Piezo1 in these neurons. However, there is no direct evidence supporting this yet. Nevertheless, with their relatively high expression of Piezo1, ultrasound could provide a way to modulate these neurons to study relevant questions in neuroscience.

## Supporting information

S1-1 Stacked slices of brain slices staining with Piezo1 in Ctrl mouse model.

S1-2 Stacked slices of brain slices staining with Piezo1 in P1KO mouse model.

## Acknowledgements

This work was supported by Guangdong High Level Innovation Research Institute (2021B0909050004), the Hong Kong Research Grants Council General Research Fund (15104520, 15102417 and 15326416), Hong Kong Innovation Technology Fund (MRP/018/18X and MHP/014/19), Shenzhen-Hong Kong Macaus Science and Technology Program (Category C), Key-Area Research and Development Program of Guangdong Province (2018B030331001), internal funding from the Hong Kong Polytechnic University Research Institute of Smart Ageing (1-CD76) and Hong Kong Polytechnic University (1-ZE1K, 1-BBAU, and 1-ZVW8). The authors would like to thank the facility and technical support from University Research Facility in Life Sciences (ULS) and University Research Facility in Behavioral and Systems Neuroscience (UBSN) of The Hong Kong Polytechnic University.

## Methods

### Mice

Animal use and care were performed following the guideline of the Department of Health - Animals (Control of Experiments) of the Hong Kong S.A.R. government. Mice were housed under standard housing condition with 12-hour light/dark cycle food and water available *ad libitum*. Transgenic mice of the lines C57BL6/J (JAX# 000664) and Piezo1^tm2.1Apat^ (JAX# 029213) were maintained for experiments.

### Stereotaxic surgery

Male, 4-8 weeks Piezo1^tm2.1Apat^ mice were deeply anesthetized by 100 mg/ml ketamine and 10 mg/ml xylazine in 0.9% NaCl and positioned in a stereotaxic injection frame (RWD Ltd, China). The anaesthetized mice were positioned in the stereotaxic apparatus, and ointment was applied on the eyes. Skin incisions were then performed to expose the skull. 400nl of virus was delivered to 3 locations respectively by a microinjection system (Nanoliter 2010, WPI Ltd) through a craniotomy (<1 mm^2^). The injection was performed unilaterally to the right motor cortex (at mm): AP: 0.0, ML: -1.0; AP: +0.5, ML: -1.0; AP: +0.5, ML: -1.5 with DV: -1. Mice injected with AAV2/9-Syn-mCherry and AAV2/9-Syn-Cre-mCherry were ‘Ctrl’ and ‘P1KO’ mice respectively, prepared for behavior, EMG and c-Fos experiments. Some mice were also simultaneously injected with AAV2/9-Syn -GCaMP6s to prepare Ctrl and P1KO mice for calcium imaging and FP recordings. The final VG/ml of all viruses was 5 × 10^12^ VG/ml (all virus from OBiO Technology, Shanghai). There are also 24 mice randomly dived into 4 groups injected with 0.2 µL rAAV-hSyn-GCaMP6s + 0.2 µL pAOV-hSyn-mCherry-2A-Cre / hSyn-mCherry / GFAP-Cre-mCherry / GFAP-mCherry into the CeA area. The coordinates used for the CeA region were AP: -1.22 mm, ML: -2.9 mm, DV: -4.60 mm.

For mice prepared for the fiber photometry experiments, after finishing the injection, an optic fiber was implanted. The fiber was fixed to the skull with glue and dental cement and allowed to set for 20 min. Scalp tissue was disinfected, and the mice were moved back to their original housing areas.

### Genotyping and tissue isolation

The success of conditional knockout was determined by PCR analysis of DNA extracted from brain tissue where the virus was injected. Genotyping was performed according to the protocol as previously specified (Cahalan *et al*., 2015). The following primers were used as indicated in Fig 1B: P1 F: GCC TAG ATT CAC CTG GCT TC; R: GCT CTT AAC CAT TGA GCC ATC T; P1 KO F: CTT GAC CTG TCC CCT TCC CCA TCA AG; R: AGG TTG CAG GGT GGC ATG GCT CTT TTT and Phire II polymerase (Thermo-Scientific #F-170S). PCR was running following the cycling conditions: Initial denaturation 98°C for 5 mins, followed by 98°C for 5 s, then 10cycles of 65℃(−0.5°C/cycle) for 5 s, 68℃for 20s, followed by 30 cycles of 98°C for 5 s, 60°C for 5 s, 72°C for 20 s, followed by a final hold of 72°C for 1 min. Reactions were separated on 2% agarose gels, yielding the following band sizes: WT: 188 bp, P1: 380 bp, P1KO: 230 bp.

### Calcium imaging *ex vivo*

Mice were deeply anesthetized by intraperitoneal injection of Ketamine/Xylazine mixture (100 mg/kg and 10 mg/kg body weight) and transcranially perfused with NMDG aCSF containing 92mM NMDG, 2.5mM KCl, 1.25mM NaH_2_PO_4_, 20mM NaHCO_3_, 10mM HEPES, 25mM glucose, 2mM thiourea, 5mM Na-ascorbate, 3mM Na-pyruvate, 0.5mM CaCl_2_·4H_2_O and 10mM MgSO_4_·7H_2_O and 12Mm NAC (pH titrated to 7.3–7.4 with concentrated HCl). Coronal brain sections 300-400µm thick were cut with a vibratome (Leica, VT1000S) in ice-cold NMDG aCSF. Slices were short recovered for 10-12 minutes at 32℃in NMDG aCSF. After that, slices were recovered in HEPES aCSF containing 92mM NaCl, 2.5mM KCl, 1.25mM NaH_2_PO_4_, 30mM NaHCO_3_, 20mM HEPES, 25mM glucose, 2mM thiourea, 5mM Na-ascorbate, 3mM Na-pyruvate, 2mM CaCl_2_·4H_2_O and 2mM MgSO_4_·7H_2_O at room temperature for at least 1 hour. The slice was then transferred to the recording ACSF (rACSF) perfusing confocal dish for the calcium image. The rACSF contained 126 mM NaCl, 1.6 mM KCl, 1.2mM NaH_2_PO_4_, 1.2mM MgCl_2_, 2.4mM CaCl_2_, 18mM NaHCO_3_, 11mM glucose. All aCSF solutions were saturated with carbogen (95% O_2_ /5% CO_2_) prior to use to ensure stable pH buffering and adequate oxygenation. A 0.5 MHz plane transducer was placed in a handmade tube, which allowed control of the stimulation area to 4 mm^2^ and was adjusted into a position underneath the rACSF confocal dish. Ultrasound stimuli (0.35 MPa) were administered with 500 µs tone burst, 1 kHz PRF, 300ms duration. The fluorescence change indicated by GCaMP6s fluorescence intensity, was calculated by ΔF/F_0_ = (F-F0)/F0, taking F_0_ to be the baseline of the fluorescence signal averaged over 10-50 s before the ultrasound onset in each session. Quantification and calculations for these experiments were performed by an experimenter blinded to the identity of the groups.

### Video capture of limb movements

Both video capture and EMG experiments were conducted after 4-6 weeks of viral injection. Under isoflurane/O_2_ anesthesia, the mice’s heads were shaved, and a 0.5MHz lithium niobate ultrasound transducer was coupled to the skull using ultrasound gel (Parker Aquasonic 100). For video capturing, 0.35/0.45 MPa ultrasound stimuli were administered with 500 µs tone burst, 1 kHz PRF, 50/ 250/ 500 ms duration and 5 s intervals. The experimenter performing the US stimulation was blinded to which group the animal belong to. Videos were captured at a rate of 30 Hz which was later analyzed by an open-source machine learning toolkit for animal pose estimation DeepLabCut. Forelimb and hindlimb trajectories after ultrasound stimulation were calculated. The greatest movement of forelimb and hindlimb digital ends within 3 s after ultrasound stimuli were extracted in terms of distance away from the position when corresponding ultrasound stimulus was applied. For behavioral tests in mice that were stimulated in either ipsi- or contralateral hemispheres, the experimental parameters were kept the same, expect for the ultrasound waveform being shaved smoothed through ultrasound modulator, and the pluses were triggered randomly (RIGOL, DG1032).

### EMG experiments

For EMG experiments, ultrasound stimuli (0.3, 0.35, 0.4, 0.45 MPa) were administered with 500 μs tone burst, 1 kHz PRF, 500 ms duration and 15 s intervals. EMG data were collected from the left gastrocnemius muscle through fine needle electrodes connected to a Meduda (Bio-Signal Technology). Data were recorded at 1000 Hz, pre-amplified and digitized. Using MATLAB (Mathworks, USA), digitized waveforms were filtered by 50 Hz notch filter and 300 Hz high-pass filter. RMS envelope of the filtered waveforms were extracted for EMG identification and feature extraction. After smoothing the envelope, mean amplitude of ‘quiet period’ prior any ultrasound stimulation was taken as baseline. If any peak rose above 4 times the baseline within 1 s window after US stimulation onset, whose waveform remained lower than 2 times the baseline within a 300 ms window prior to the onset of stimulation, the peak was considered a successful US-induced EMG response and the amplitude was extracted.

### Fiber Photometry

Following implantation and recovery, around 4 weeks, mice were anesthetized with isoflurane (1.0-2.5% in O_2_). Ultrasound gel was applied on a shaved head, to promote acoustic coupling. A 0.5 MHz transducer embedded with a water tube wave-guide was installed above the skull. Mice were stimulated with four trials of ultrasound stimulation (0.03 MPa, 0.06 MPa, 0.12 MPa or 0.25 MPa). Each trial was 3-6 ultrasound stimuli, with 5 s interval between each pulse. Mice were allowed to rest for 50 s between trials. GCaMP6s fluorescence was captured with a fiber photometry system (Thinker Tech Nanjing BioScience Inc.). The excitation and receiving wavelength for fiber photometry were 470 nm with 30 nm bandwidth and 510 nm with 25 nm bandwidth, respectively. Data was collected at 100 Hz and analyzed using a customized MATLAB script. The relative change in fluorescence, ΔF/F_0_ = (F-F0)/F0, was calculated by taking F_0_ to be the baseline of the fluorescence signal averaged over 1 s before the ultrasound onset in each session.

### Immunohistochemical fluorescent staining

Mice were sacrificed 90 minutes after ultrasound treatment and perfused with PBS, followed by 4% paraformaldehyde (PFA) (cat. no. P1110, Solarbio) in PBS. After dissection, brains were post-fixed overnight in 4% PFA and then rinsed in PBS. Starting from the injection plane, around 10 continuous coronal brain slices with the interval of 105 μm were collected. Slices were blocked using blocking medium containing 10% normal goat serum, 1% BSA and 0.3% Triton X-100, for 90 mins and incubated overnight in primary antibody solution diluted in blocking. Slices were then washed and incubated with secondary antibodies diluted in PBS for 90 minutes at room temperature. Slices were then washed, stuck to glass slides, dried, and mounted on coverslips using small drops of Prolong Diamond Antifade Mountant with DAPI and allowed to cure in the dark at room temperature overnight. Coverslip edges were sealed using transparent nail enamel and imaged using a confocal laser scanning microscope (TCS SP8, Leica). Primary antibodies used were c-Fos (2250, CST, dilution 1:500), Piezo1 (ab128245, abcam, dilution 1:50), and MAP2 (PA1-16751, Invitrogen, dilution 1:500). For the blocking experiments, specific Piezo1 immunizing peptide (ab133015, abcam, concentration: 20 μg/ml) was added to the Piezo1 antibody for 2 hrs preincubation. Secondary antibodies, used at a dilution of 1:1000, were goat anti-rabbit IgG (H+L) Alexa Fluor 488 (A-11008, Invitrogen) and goat anti-mouse IgY (H+L) Alexa Fluor 633 (A-21103, Invitrogen). The c-Fos (green) stained neurons in were 853 × 853 μm slice were counted using ImageJ. The counting was single-blinded, performed by an experimenter who did not know the groups beforehand.

### Statistics

Statistical analyses were performed in GraphPad Prism. All statistical tests in this study were two-tailed. Single-variable comparisons were made with t-test. Group comparisons were made using either analysis of variance (ANOVA) followed by Tukey post-hoc analysis, Holm-Sidak test or Dunnett’s test.

## Supplemental Material

**Vid S1**. Stacked slices of brain slices staining with Piezo1 in Ctrl and P1KO mouse model.

**Vid S2**. Video of ultrasound-induced hindlimb movement in Ctrl mice (Ultrasound stimulation to the ipsilateral side of virus injection)

**Vid S3**. Video of ultrasound-induced hindlimb movement in P1KO mice Ultrasound stimulation to the ipsilateral side of virus injection)

**Vid S4**. Video of ultrasound-induced hindlimb movement P1KO mice Ultrasound stimulation to the contralateral side of virus injection)

**Fig S1.**
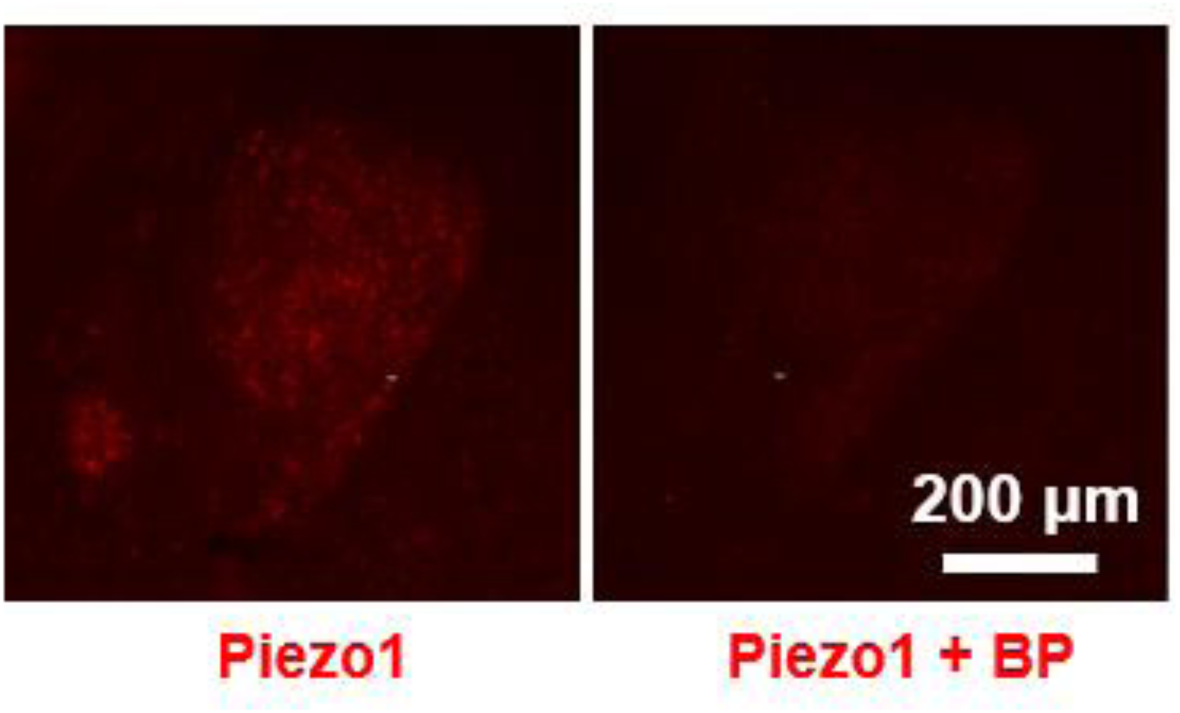
Immunostaining of Piezo1 with or without Piezo1 blocking peptide. Left: Piezo1 staining in CEA region. Right: Piezo1 staining in the CEA, which was pre-incubated with the Piezo1 antibody blocking peptide.

**Fig S2.**
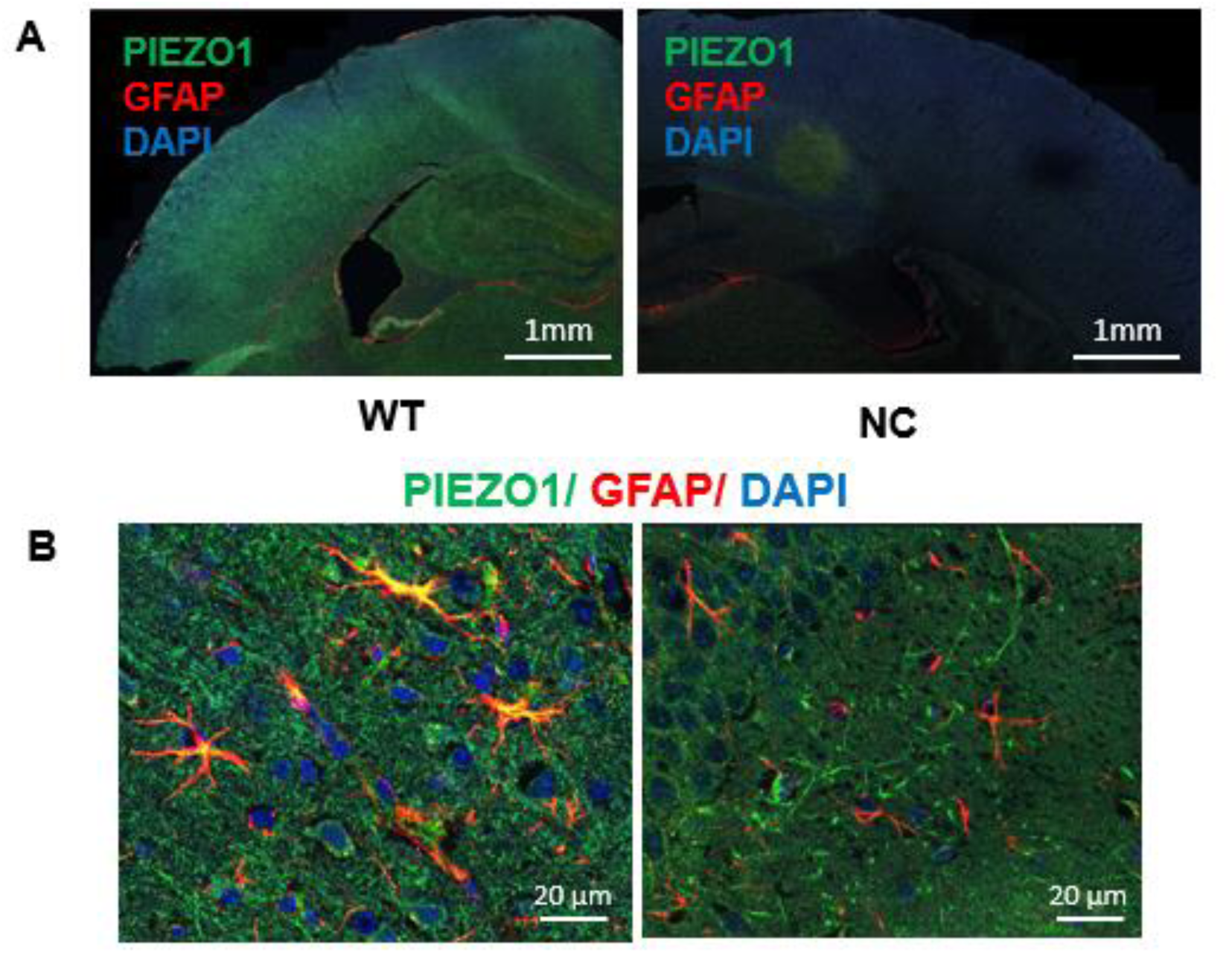
Immunostaining of Piezo1 and GFAP in mouse brain. **(A)** Left: Co-staining of Piezo1 and GFAP in the mouse brain. Right: Negative control (no primary antibody) of the co-staining of Piezo1 and GFAP in mouse brain. **(B)** Left: Higher magnification image of co-localized of Piezo1 and GFAP, indicating Piezo1 expression in some astrocytes. Right: Higher magnification image of a region with non-co-localized Piezo1 and GFAP.

**Fig S3.**
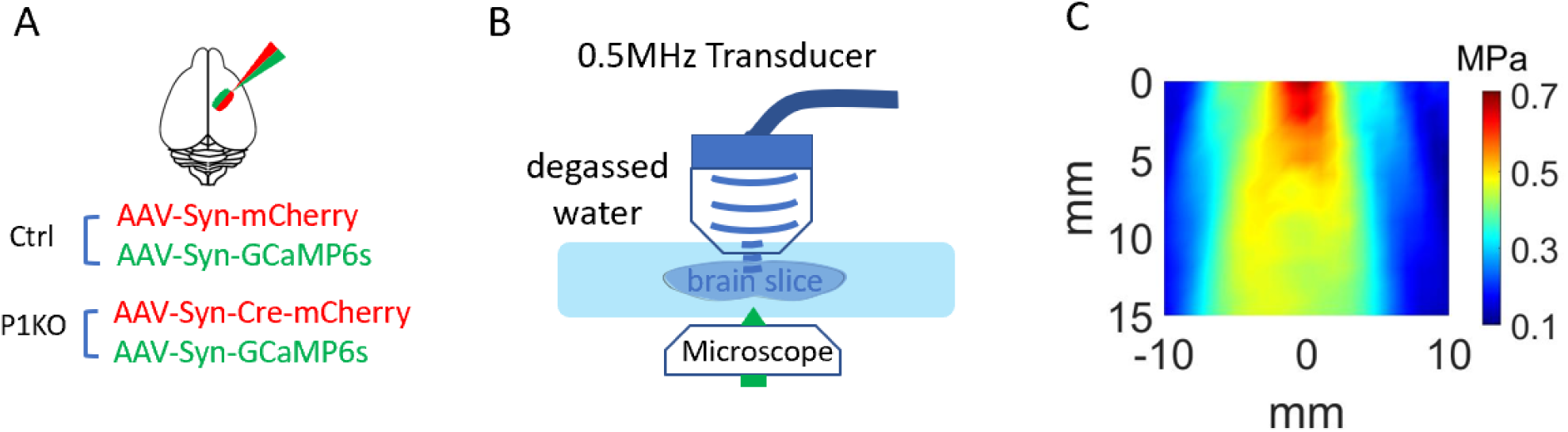
Procedure of calcium imaging under ultrasound stimulation in *ex vivo* brain slices. **(A)** Ctrl mice were prepared by dual virus injection of mCherry and GCaMP6s in cortex in P1^f/f^ mice while P1KO mice were injected with mCherry-Cre and GCaMP6s. **(B)** Schematic illustration of the setup for calcium imaging and ultrasound stimulation in *ex vivo* brain slices. **(C)** Acoustic mapping profile of the ultrasound field generated by the transducer within our imaging setup, indicating that the intensity received by the sample would be about 0.35 MPa.

**Fig S4.**
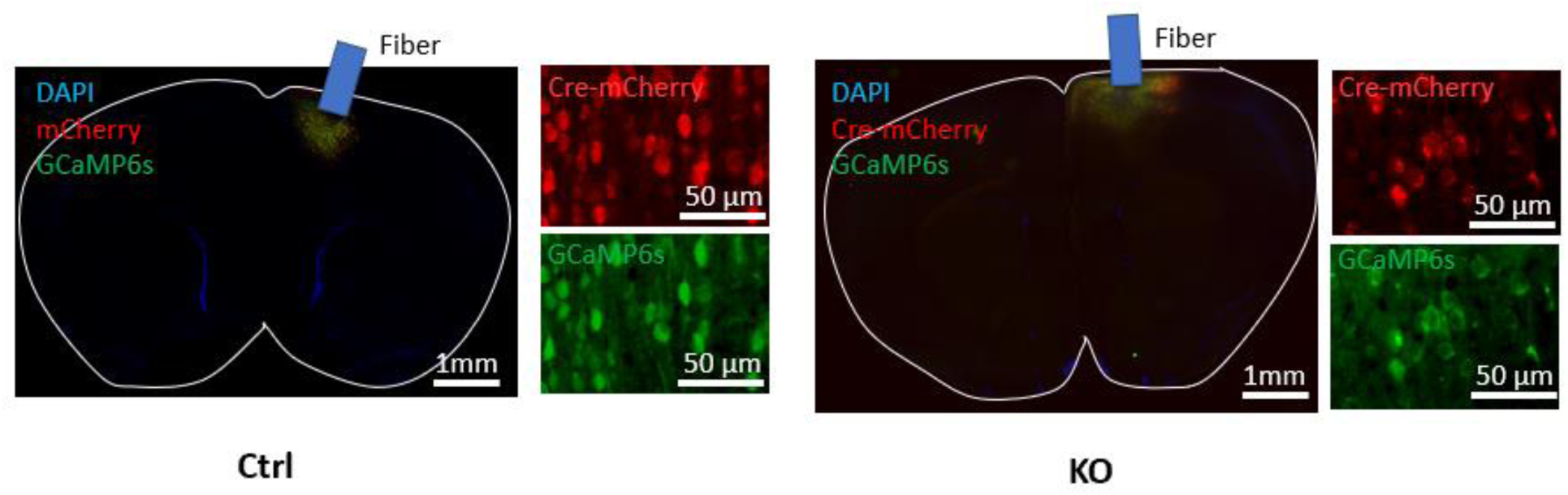
Dual expression of mCherry + GCaMP6s in Ctrl mice and mCherry-Cre + GCaMP6s in P1KO in mice used for FP. Left: Low-magnification image showing a representative image of dual virus expression in the right motor cortex with 2 high-magnification images showing a representative example of mCherry and GCaMP6s expression in Ctrl mice; Right: the same set of representative images of dual expression Cre-mCherry and GCaMP6s in the motor cortex of P1KO mice.

**Fig S5.**
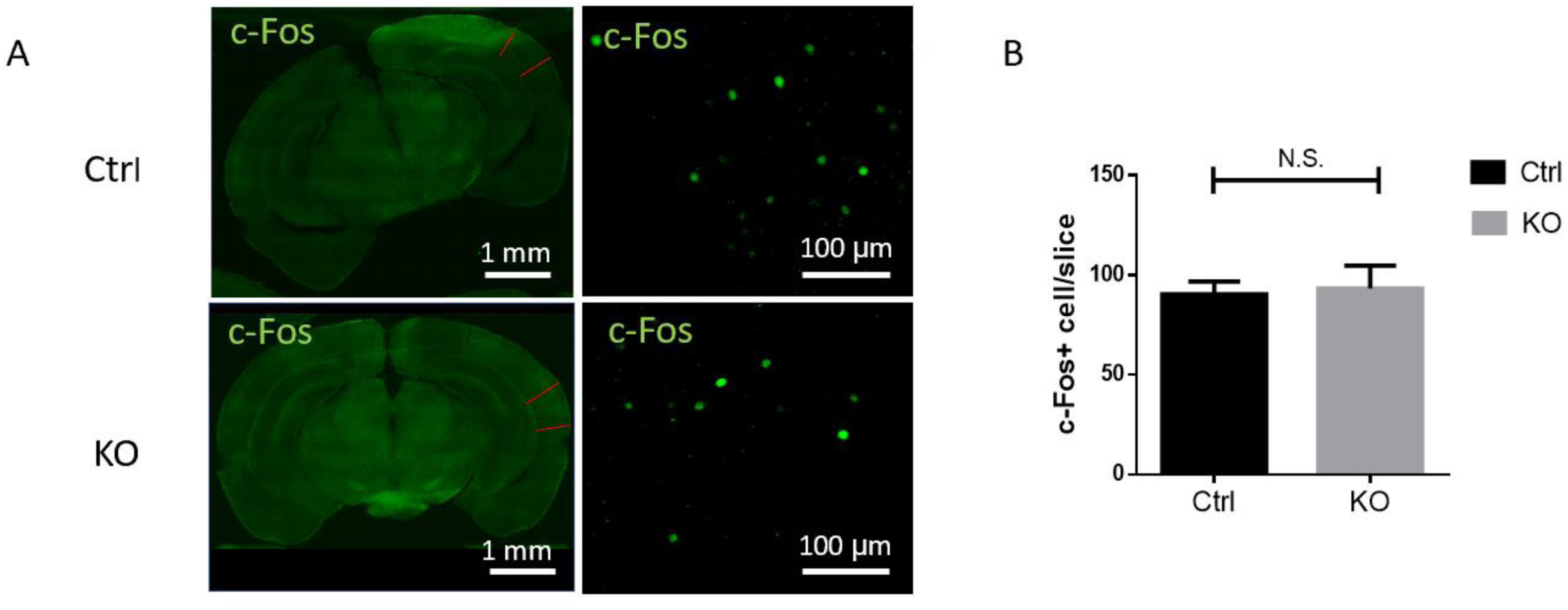
Piezo1-mediated ultrasonic neuromodulation did not trigger widespread activation of the auditory cortex. **(A)** Representative of images of c-Fos expression in the auditory cortex of Ctrl and P1KO mice after 0.45MPa ultrasound stimulation for 50 mins. Top: Representative low- and high-magnification images showing c-Fos expression in Ctrl mice; Bottom: The same representative images in P1KO mice; Green: c-Fos. **(B)** Summarized c-Fos number per slice. n = 11 slices each from 3 P1KO and 3 Ctrl mice. Bars represent mean ± SEM, ‘N.S.’ not significant, by unpaired t-test.

**Fig S6.**
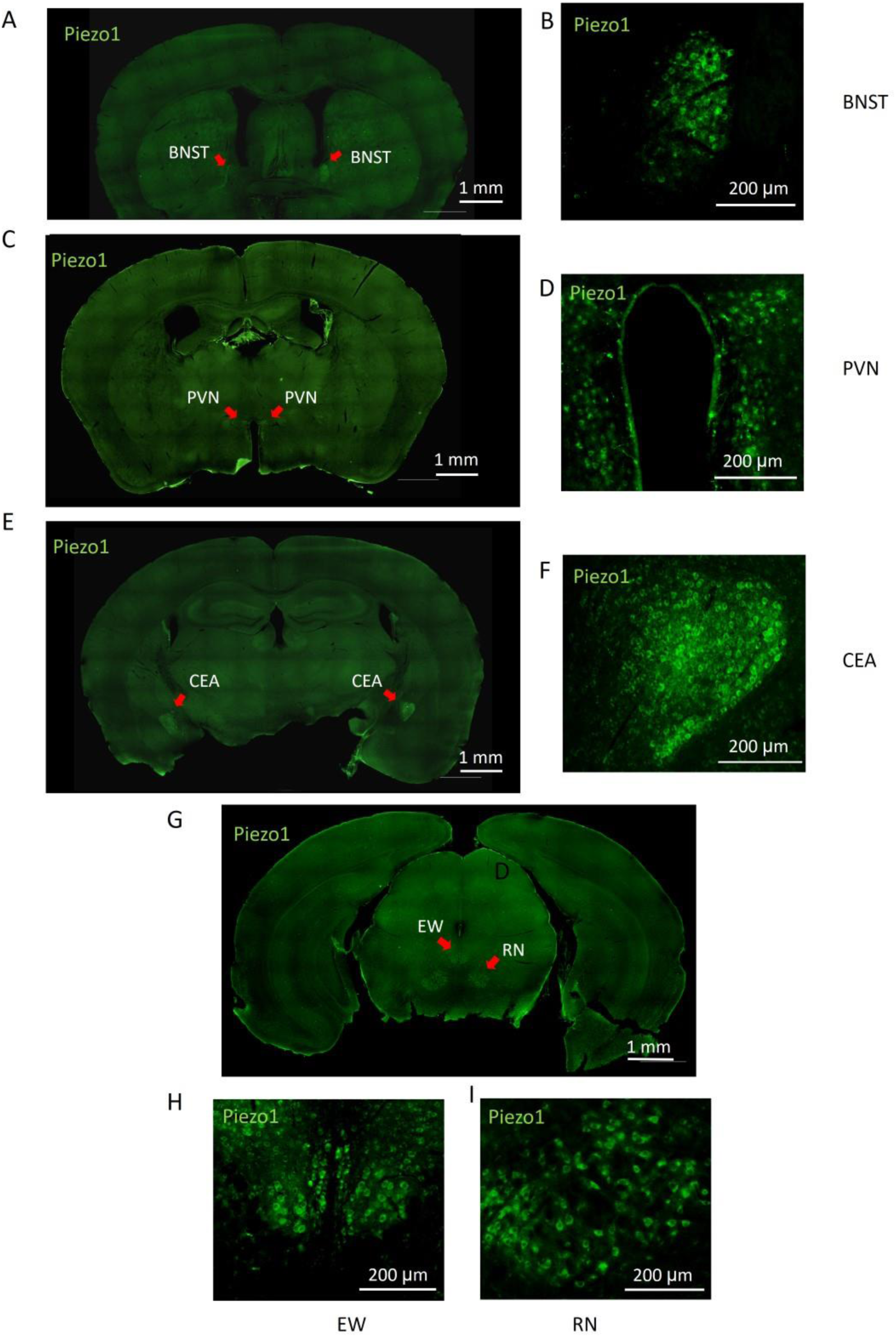
Immunostaining of Piezo1 in different brain regions. **(A, C, E, G)** low-magnification images of Piezo1 stained in BNST, PVN, CEA, EW and RN. Green: Piezo1.I**(B, D, F, H, I)** high-magnification images of Piezo1 stained in BNST, PVN, CEA, EW and RN. Green: Piezo1.I

**Fig S7.**
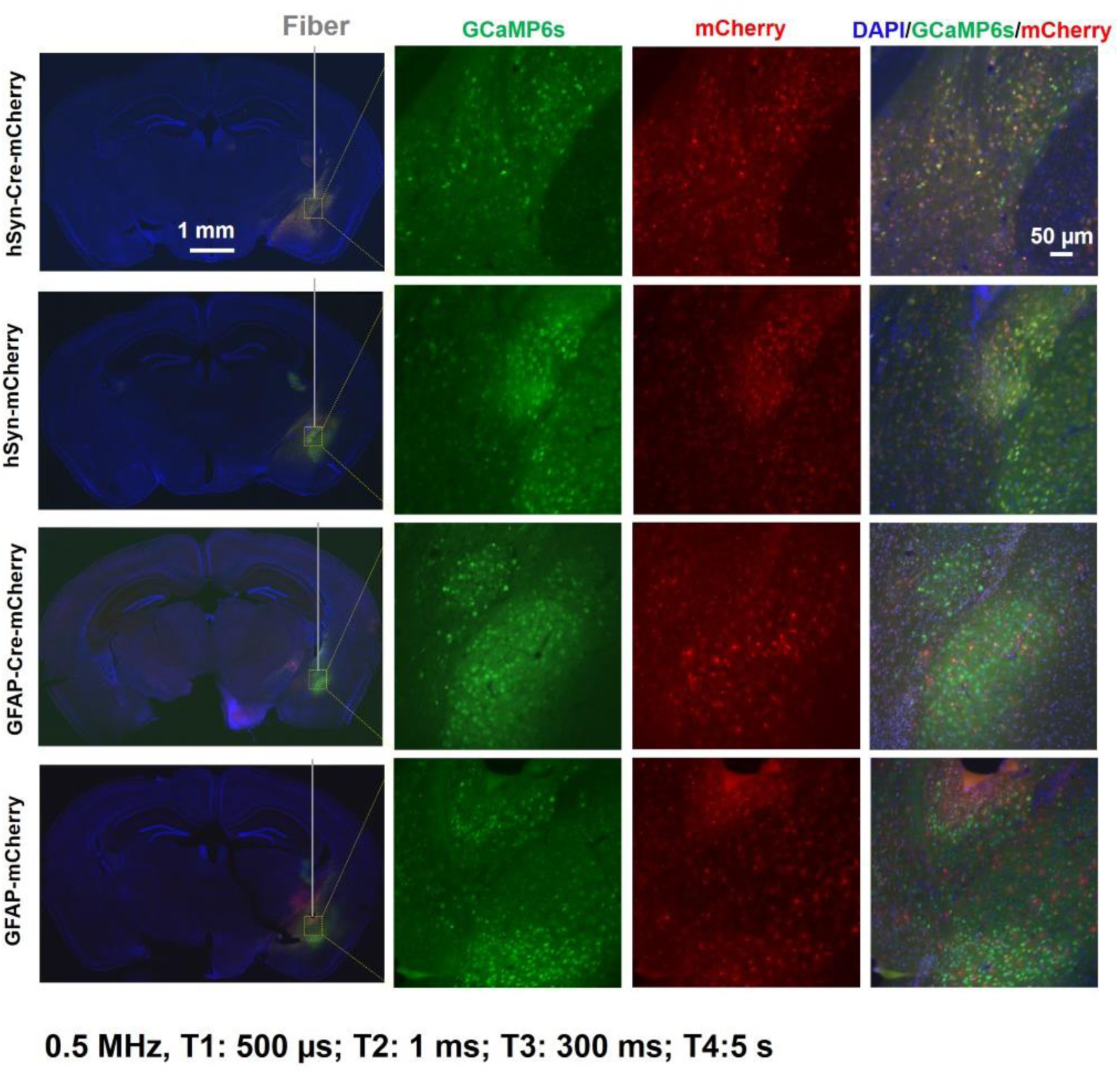
Dual expression of mCherry and GCaMP6s in mice in hSyn, hSyn-Cre, GFAP, GFAP-Cre groups (from up to bottom) in FP experiment. Left: Representative low-magnification images showing virus expression in the right CEA region. Right: 3 high-magnification images of the same.

